# DAQplugin: Interactive Deep Learning-Based Validation of Cryo-EM Protein Models in ChimeraX

**DOI:** 10.64898/2026.06.11.731735

**Authors:** Genki Terashi, Han Zhu, Daisuke Kihara

## Abstract

Although an increasing number of protein structures are determined by cryogenic electron microscopy (cryo-EM), structure modeling frequently suffers from residue misassignments and sequence register shifts, particularly in regions with ambiguous density. Here, we present DAQplugin, a ChimeraX plugin for real-time evaluation of protein models against cryo-EM density maps using the deep-learning-based residue-wise model quality (DAQ) score. Unlike existing validation tools that are typically applied after model construction, DAQplugin enables interactive validation during model building and refinement. DAQ has been shown to accurately identify residue assignment errors, including sequence register shifts, as well as local conformational modeling errors. DAQplugin also provides guidance for correcting sequence register shifts by suggesting alternative residue placements along the backbone. The plugin is computationally efficient and runs on standard CPUs without requiring GPU hardware, enabling deep-learning-based validation on ordinary laptops during interactive model building, model–map fitting, and refinement. DAQplugin facilitates more accurate interpretation of cryo-EM density maps and improve the reliability assessment of protein structure models.

**Highlights:**

- DAQplugin enables real-time deep-learning validation of cryo-EM models in ChimeraX
- DAQplugin detects sequence-register and positional errors and guides interactive refinement
- Precomputed DAQ grids enable rapid model–map evaluation on standard CPUs

## 1. Introduction

Protein structure determination using cryogenic electron microscopy (cryo-EM) has expanded rapidly in recent years, enabling structural characterization of increasingly large and complex macromolecular assemblies^1^. Despite these advances, accurate model building remains challenging, particularly in regions with ambiguous or low-resolution density, where residue misassignments and sequence register shifts are often observed.

Therefore, structural validation has become an urgent task as the community has noted^2^. Several methods have been developed to validate protein models built into cryo-EM maps^3^. Existing validation metrics generally fall into two categories: map-model scores and model-coordinate scores. Map-model scores, such as correlation coefficients and Q-scores^4^, evaluate the agreement between an atomic model and the density map. But these methods often fail to detect residue misassignments because incorrect side-chain identities or sequence register shifts can still produce good local density agreement when the backbone is correctly positioned. On the other hand, model-coordinate scores, including Ramachandran analysis and clash detection^5^, assess stereochemical quality but do not directly evaluate consistency with the cryo-EM density.

We previously developed the deep-learning-based residue-wise model quality (DAQ) score^6^. DAQ evaluates the likelihood that each residue assignment is consistent with local cryo-EM density by recognizing density patterns corresponding to amino acid types, secondary structures, and Cα atom positions. Consequently, DAQ is able to identify residue assignment errors, including sequence register shifts and local conformational errors, that are often not easy to detect using conventional validation metrics.

DAQ is available to the structural biology community as stand-alone software or on a server platform^7^. In addition, evaluation results of existing PDB entries are available at the DAQ-score database^8^ and the Protein Data Bank Japan^9,10^. However, the original implementation was designed for post hoc validation after model construction. Applying DAQ during interactive model building requires recomputing the scores each time the model coordinates are modified, interrupting the modeling workflow and preventing real-time feedback during model building and refinement.

This limitation is common to essentially all existing cryo-EM model validation tools, which are designed for post hoc rather than interactive use. Although widely-used modeling environments such as Coot^11^ and UCSF ChimeraX^12^ support interactive model building and refinement through tools such as Refmac^13^ and ISOLDE^14^, validation metrics, including MolProbity^5^, Ramachandran analysis, atom-clash detection for stereochemical property evaluation, as well as map-model correlation, and Q-score^4^ for cryo-EM map and model fit evaluation, must be recomputed after model modifications. Consequently, error detection and model correction are separated into iterative validation–refinement cycles, reducing the efficiency of interactive model building.

Here, we present DAQplugin, a ChimeraX plugin for real-time evaluation of protein models against cryo-EM density maps using the DAQ framework. DAQplugin computes DAQ scores directly within ChimeraX and interactively displays residue-level model–map agreement in real-time as the model is modified. For potential sequence register shifts, the plugin also indicates the suggested direction of sequence reassignment using graphical arrows, enabling users to identify and correct modeling errors immediately during structure construction. When users modify a structure interactively, atomic positions can be dynamically refined using molecular dynamics in ISOLDE ^14^. Because ISOLDE occasionally generates rare side-chain rotamers, we also developed DaQRotFix (Density-assisted Quick Rotamer Fix), a tool for correcting rotamer outliers.

Although the original DAQ framework relies on GPU-based deep learning, DAQplugin has been redesigned to run efficiently on standard CPUs, allowing installation and use on ordinary laptops without specialized hardware or complex software dependencies. This efficiency is achieved by restructuring the algorithm so that the computationally intensive 3D convolutional neural network inference is performed only once during initialization. To our knowledge, DAQplugin is the first tool to provide interactive deep-learning-based validation of cryo-EM protein models within a molecular graphics environment. By integrating residue-level validation directly into the model-building workflow, DAQplugin transforms model validation from a post hoc procedure into an integral component of interactive structure determination.

## 2. Methods

### 2.1. Overview of the DAQ

First, we discuss how DAQ is computed with an emphasis on the difference between the original DAQ computation and this plugin version. For details of the algorithm and benchmark results, refer to the original paper^6^. The overall workflow of DAQplugin is shown in Fig. 1a. The method consists of two main components: (1) precomputation of DAQ scores and (2) real-time scoring and visualization.

**Figure 1.**
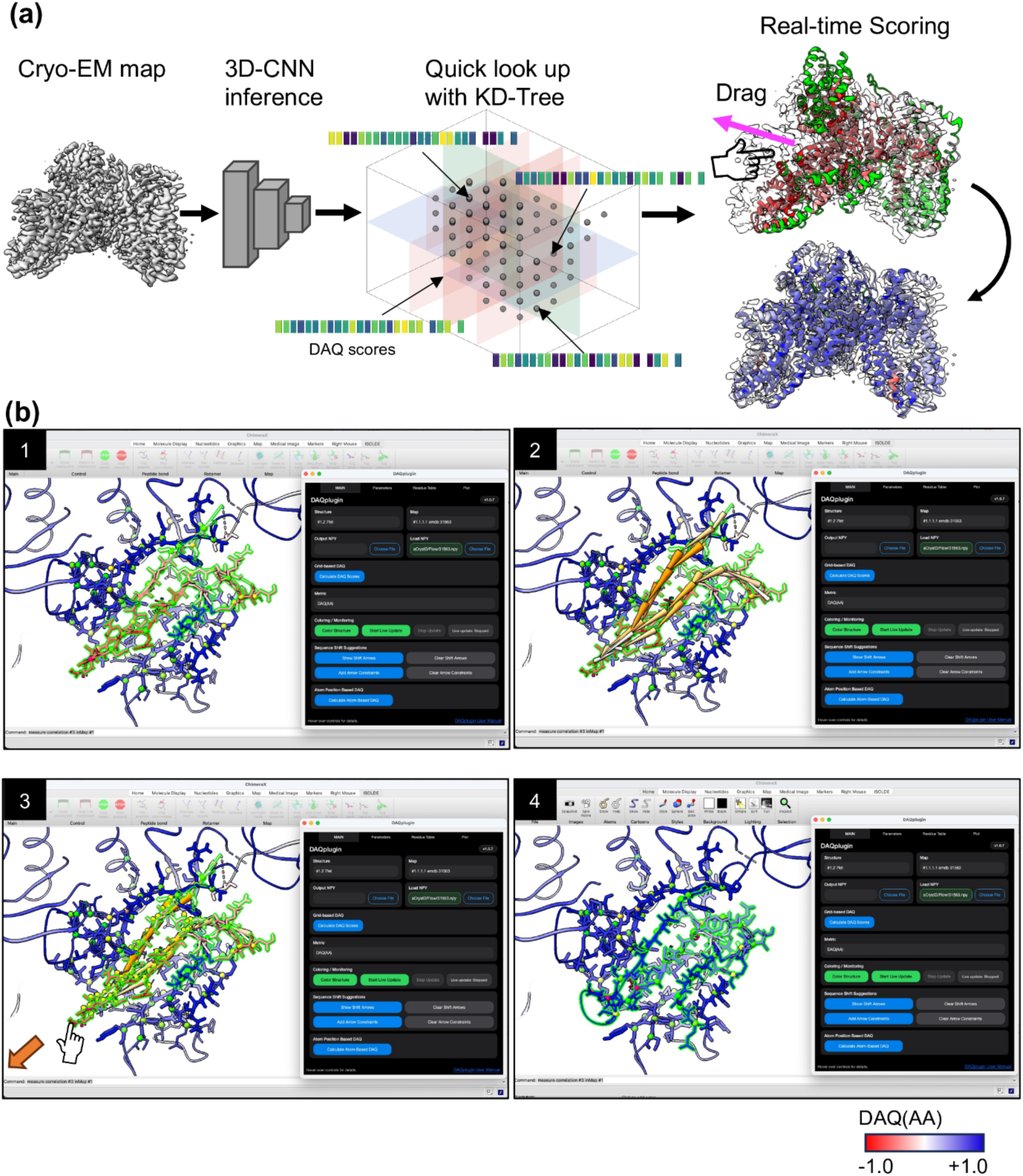
Overview of the DAQplugin workflow and GUI. (a) A cryo-EM density map (EMD-9906) is processed by a 3D convolutional neural network (3D-CNN) to predict probabilities for 20 amino-acid types, Cα atoms, and three secondary-structure classes. These predictions are converted into precomputed DAQ score grids [DAQ(AA), DAQ(Cα), and DAQ(SS)] and indexed using a KD-tree. The translucent planes illustrate the recursive spatial partitioning of the KD-tree, enabling rapid retrieval of DAQ scores near atomic coordinates. During interactive model building and refinement in ChimeraX, scores are retrieved from the precomputed grids and mapped onto the atomic model, allowing DAQ visualization to update in real time as the model is modified. (b) Example of real-time DAQ score updating during interactive refinement of PDB 7FET chain B with the EM map (EMD- 31563). ChimeraX’s GUI interface and DAQplugin GUI are shown for each screenshot. The computation was also recorded in Movie S1. (1) DAQplugin identifies a low DAQ(AA) score region (Gly593-Pro621, average DAQ(AA) score −0.51). (2) DAQplugin computes the suggested sequence register shift, shown as orange arrows. The intensity of the orange arrows indicates the magnitude of the predicted improvement in the DAQ(AA) score. (3) Using ISOLDE, the user manually drags that region toward the left. Position constraints (yellow pins) also guide the simulation. DAQ scores are updated in real time during the refinement process. (4) After the refinement, the modified region shows improved DAQ scores (average 0.89).

DAQ evaluates the consistency between a protein structure model and a cryo-EM density map at the residue level. It employs a three-dimensional convolutional neural network (3D-CNN) trained to recognize structural features, including amino acid types, Cα atom positions, and secondary structures, from local density patches (11 × 11 × 11 voxels). The input density patch is sufficiently large to capture not only the density of the target residue but also the surrounding structural environment, including neighboring residues that interact with it. The network predicts residue-wise probabilities for amino acid identity, Cα position, and secondary structure, which are subsequently used to compute the DAQ score.

The input to DAQ is a cryo-EM map of up to 5.0 Å global resolution and a protein structure model built from the map. The map is first resampled to the grid size of 1.0 Å if it has a different grid size. Then, the 3D-CNN processes local density patches and computes the probability of the presence of twenty amino acid types, Ca atoms, and three protein secondary structures at each grid position of the map.

For amino acid type assessment, the network computes a probability array

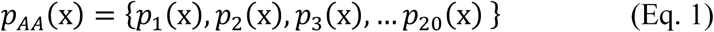

where *p_i_*(x) represents the computed probability that the position x corresponds to amino acid type *i* among the 20 standard amino acids. The DAQ score of residue *r* at a grid position is defined as a log-likelihood ratio:

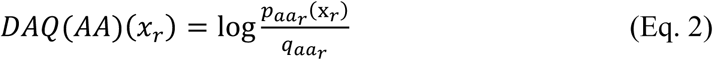

where *aa_r_* denotes the amino acid type assigned to residue *r* at position *x_r_*, and *q*(*aa_r_*) is the background probability of amino acid *aa_r_*, computed as the average predicted probability of that amino acid over the entire density map. A positive DAQ(AA) score indicates that the local density is more consistent with the assigned amino acid type than expected from the background distribution, whereas a negative score indicates lower-than-expected consistency. Therefore, large negative DAQ(AA) scores suggest incorrect residue assignment or inconsistency between the model and the local cryo-EM density. In the original DAQ study, we showed that a 19-residue moving-average DAQ(AA) score below −0.5 is a strong indicator of sequence assignment errors^6^.

The DAQ score of the C*a* atom *DAQ*(*Ca*)(*x_r_*), is computed as

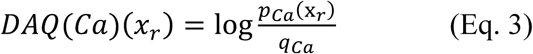

where *p_Ca_*(x*_r_*) is the probability predicted by the 3D-CNN that the local density around position x*r* corresponds to the backbone Cα atom, and *q_Ca_* is the corresponding background probability. DAQ(Cα) evaluates whether the local density at the modeled Cα atom position is consistent with the density patterns expected for protein Cα atom. Positive DAQ(Cα) scores indicate higher-than-background consistency, whereas negative scores indicate lower-than-background consistency, suggesting potential errors in backbone positioning.

DAQ score is also provided for the secondary structure. *DAQ*(*SS*)(*x_r_*), is computed as

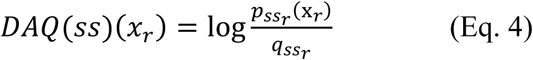

where *ss_r_* denotes the secondary structure assigned to residue *r* by DSSP^15^. *p_ss_*_r_(x*_r_*) is the probability predicted by the 3D-CNN that the local density around position x*r* corresponds to secondary structure *ssr* and *q_ss_*_r_ is its background probability. In the original DAQ implementation, secondary structure information was obtained from SPOT-1D^16^ sequence-based predictions, with the three secondary structure classes weighted by their predicted probabilities. In DAQplugin, we instead assign secondary structures directly from the current atomic model using DSSP for more direct comparison between the modeled secondary structure and the local cryo-EM density.

### 2.2. Real-time DAQ scoring in DAQplugin

A major challenge in applying deep-learning-based validation during interactive refinement is the computational cost of neural network inference. Recomputing the 3D-CNN predictions after every coordinate update would make real-time validation impractical.

To address this, DAQplugin performs the 3D-CNN inference only once during initialization and stores the predicted probabilities as precomputed three-dimensional grids. During model editing, residue-wise DAQ scores are obtained by quick lookup of these grids using KD-tree structure^17^, enabling real-time feedback. During initialization (Fig. 1a), the 3D-CNN is applied to grid points sampled every 2 Å across the cryo-EM map as default, and the predicted probabilities for amino acid types, Cα atoms, and secondary structures are stored. To further reduce computation, calculations are restricted to regions with density values above half of the user-defined contour level.

In the original DAQ implementation, background probabilities were computed by averaging predicted probabilities at the atomic positions of the input model. In DAQplugin, the background is instead computed once by averaging over all sampled grid points above the contour threshold, eliminating the need to recompute background probabilities whenever the model is modified. Consequently, users can immediately observe whether local model edits improve or worsen agreement with the cryo-EM density during interactive refinement.

### 2.3. Suggesting Sequence Register Shifts

Sequence register shifts, in which the amino acid sequence is misassigned while the protein backbone is correctly modeled, are among the most common errors in cryo-EM protein structure models^6^. Because DAQ evaluates residue assignment directly from the cryo-EM density, it is particularly effective at identifying such errors.

DAQplugin further assists model correction by automatically suggesting the direction of sequence reassignment. When the user activates the Suggest Register Shift function, the plugin scans all residues in the protein model, evaluates alternative sequence assignments within a local region, and indicates the recommended shift direction using arrows displayed along the protein backbone.

Given a user-defined half-window size *n*(default: 5) and maximum shift distance *m*(default: 5), DAQplugin considers the residue window from *i* − *n* to *i* + *n* centered at residue *i*. For each possible sequence shift *s* (−*m* ≤ *s* ≤ *m*), the average DAQ(AA) score is computed as

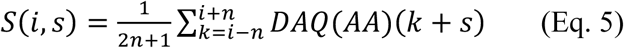

where DAQ(AA)(*k* + *s*) is calculated using the Cα position of residue *k* + *s* and the amino acid identity assigned to residue *k*. The shift producing the largest improvement relative to the original assignment (*s* = 0) is selected as the suggested register shift.

The recommended shift is visualized in ChimeraX as an arrow along the protein backbone (Fig. 1b), indicating the direction in which the sequence assignment should be moved to improve the DAQ(AA) score. Users can then adjust the sequence assignment interactively while monitoring the updated DAQ scores in real time.

Fig. 1b illustrates an example of correcting a sequence register shift using DAQplugin. A continuous stretch of negative DAQ(AA) scores (residues Gly593-Pro621 in Chain B) was detected in two β-strands of the structure model of SARS-CoV-2 B1.1.7 Spike Glycoprotein (PDB ID: 7FET; EMDB: 31563), indicating a likely sequence register error (panel 1). When the user selected a residue within the erroneous region and activated the Suggest Register Shift function, DAQplugin indicated the direction of the optimal sequence shift with orange arrows (panel 2). Following this suggestion, the sequence assignment was interactively shifted along the backbone while maintaining stereochemical integrity using molecular dynamics refinement in ISOLDE^14^, where interatomic distances were constrained during the refinement process (panel 3). The sequence register shift correction and real-time updating of DAQ score are demonstrated in Movie S1. After applying the suggested two-residue register shift, the negative DAQ(AA) scores disappeared and were replaced by uniformly positive scores (panel 4), indicating substantially improved agreement between the protein model and the cryo-EM density.

### 2.4. Side-chain rotamer refinement using DaQRotFix

Occasionally, ISOLDE refinement produces side-chain conformations that are classified as rotamer outliers. To address this issue, we developed DaQRotFix (Density-assisted Quick Rotamer Fix) (https://cxtoolshed.rbvi.ucsf.edu/apps/chimeraxdaqrotfix), a ChimeraX plugin that corrects side-chain rotamers through local molecular-dynamics flexible fitting (MDFF)^18^ in ISOLDE.

DaQRotFix first identifies rotamer outliers using ISOLDE’s side-chain validation, which evaluates conformations against the MolProbity rotamer library^19^. By default in ISOLDE, residues with a rotamer probability below 0.05% are classified as outliers. For each identified outlier, DaQRotFix attempts to replace the original rotamer with a higher-probability canonical rotamer, testing candidates in decreasing order of probability. Each candidate rotamer undergoes local MDFF followed by energy minimization, while backbone atoms (N, Cα, C, and O) are positionally restrained. A candidate is accepted if it resolves the target rotamer outlier without introducing new outliers in neighboring residues. The first candidate satisfying these criteria is retained; if none succeeds, the original conformation is restored. We recommend running DaQRotFix after interactive ISOLDE refinement to correct any remaining side-chain rotamer outliers.

## 3. Results

### 3.1. Comparison with the grid-based DAQ and the original DAQ

The original DAQ computes background probabilities (the denominators in Eqs. 2–4) using Cα positions in the model^6^. In DAQplugin, background probabilities are instead computed over grid points within half of the user-specified contour level, avoiding the need to recompute them whenever the model is modified. We therefore examined whether DAQ scores computed using these two approaches are consistent.

Following the original DAQ study^6^, we collected PDB entries with multiple model versions derived from the same cryo-EM map. Residues in the initial model were classified as either unchanged or revised in a later deposited version. Revised residues were considered likely to contain errors in the initial model and were used to evaluate the ability of DAQ(AA) and DAQ(Cα) to distinguish potentially incorrect regions. DAQ(SS) was excluded because it is less directly related to model revision and has lower sensitivity for detecting modeling errors.

Models were obtained from the DAQ database (version 2025-0430)^8^, which contains 275,728 protein chains associated with EMDB maps. All entries in the database had already been filtered to include structures with reported resolutions of 2.5–5.0 Å and sufficient global model–map agreement (sum of cross-correlation coefficient and map-overlap score ≥1.50). From this dataset, we identified 91 protein chains with at least two deposited versions differing by >1.0 Å Cα RMSD. The distributions of DAQ(AA) and DAQ(Cα) scores were then compared between these two groups.

Figures 2a–c compare DAQ(AA) scores computed using the two background definitions. In the 91 protein chains, residues displaced by >2.0 Å between the initial and revised models, i.e. more than half the typical distance between adjacent Cα atoms, were classified as revised, yielding 8,576 revised and 36,578 unchanged residues. The grid-based DAQ(AA) implemented in DAQplugin (Fig. 2a) clearly distinguished revised (red) from unchanged (blue) residues, consistent with previous DAQ studies^6,8^. Below the −0.5 cutoff, 94.6% of residues were revised. A similar separation was observed with the original atom-coordinate-based DAQ(AA) (Fig. 2b), for which 94.1% of residues below −0.5 were revised. The two DAQ(AA) scores were highly correlated (r = 0.96; Fig. 2c), demonstrating that the grid-based implementation closely reproduces the original DAQ(AA) evaluation.

**Figure 2.**
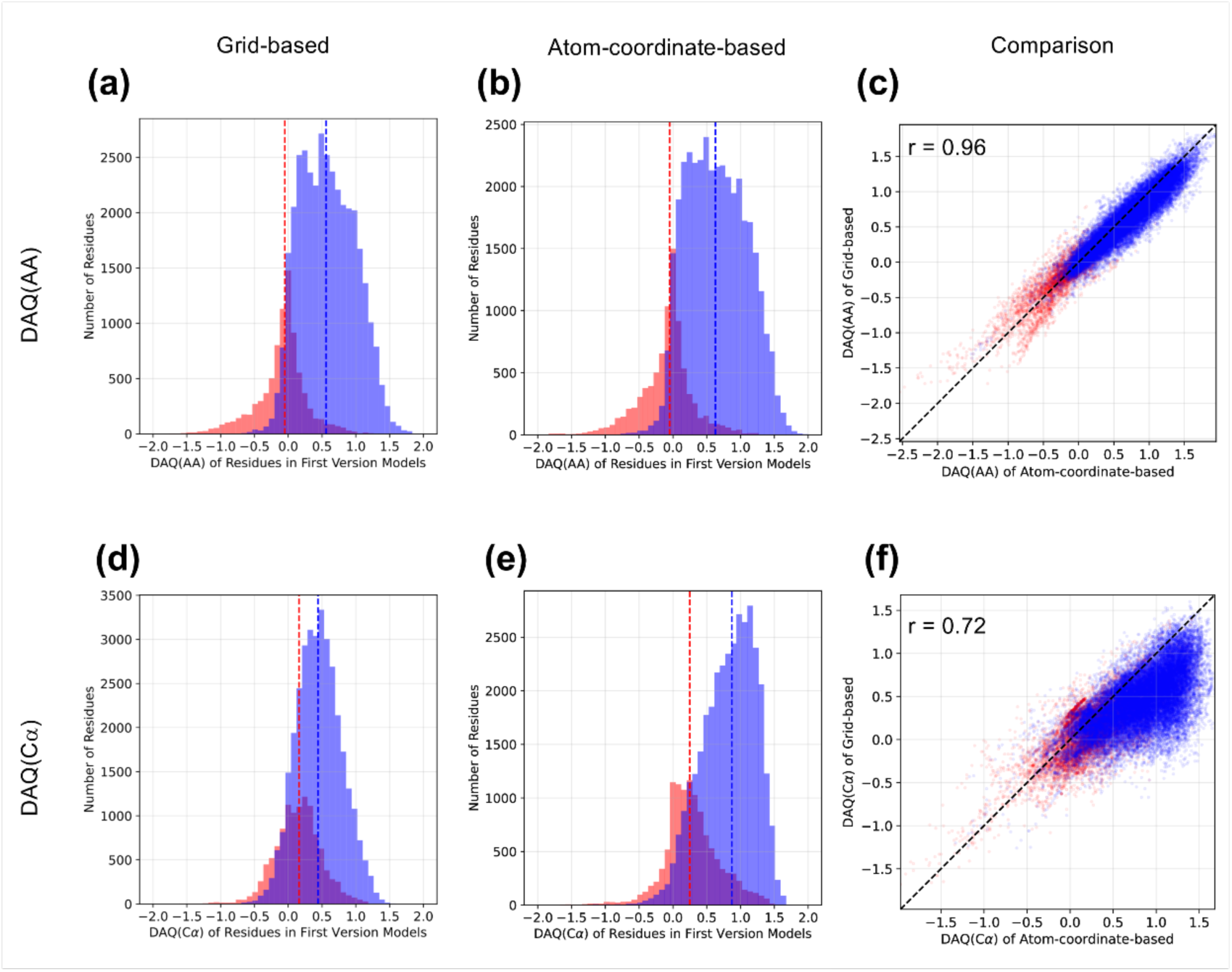
Distribution of grid-based and atom-coordinate-based DAQ scores DAQ scores were computed for first-version models of protein chains with multiple deposited versions derived from the same cryo-EM maps. Residues were classified as modified in the revised models (red) or preserved between versions (blue). Dashed lines indicate median values. **(a–c)** DAQ(AA) comparison. **(a)** Grid- based DAQ(AA) distributions for modified (>2.0 Å displacement; 8,576 residues) and preserved residues (36,578), with median scores of −0.05 and 0.56, respectively. **(b)** Corresponding atom- coordinate-based DAQ(AA) distributions, with medians of −0.05 and 0.63. **(c)** Comparison of grid- and atom-coordinate-based DAQ(AA) scores (Pearson *r* = 0.96). **(d–f)** DAQ(Cα) comparison. **(d)** Grid-based DAQ(Cα) distributions for modified (>1.0 Å displacement; 11,478 Cα atoms) and preserved Cα atoms (34,074), with medians of 0.16 and 0.44, respectively. **(e)** Corresponding atom- coordinate-based DAQ(Cα) distributions, with medians of 0.25 and 0.87. **(f)** Comparison of grid- and atom-coordinate-based DAQ(Cα) scores (Pearson *r* = 0.72).

Figures 2d–f evaluate the same models using DAQ(Cα). Cα atoms displaced by >1.0 Å between the initial and revised models—more than half the typical distance between adjacent heavy atoms—were classified as revised, yielding 11,478 revised and 34,074 unchanged Cα atoms. Although the separation between revised (red) and unchanged (blue) atoms was less distinct than for DAQ(AA), the grid-based (Fig. 2d) and atom-coordinate- based (Fig. 2e) DAQ(Cα) scores showed similar distributions. Among Cα atoms with scores ≤−0.5, 79.8% and 83.3% were revised for the grid-based and atom-coordinate-based implementations, respectively. The two DAQ(Cα) scores also showed a strong correlation (r = 0.72; Fig. 2f), supporting the consistency of the grid-based implementation with the original DAQ.

In Supplementary Fig. S1, we evaluated finer 1 Å grid sampling. The DAQ score distributions remained consistent with those obtained using the recommended 2 Å grid, while correlations with atom-coordinate-based scores increased from 0.96 to 0.99 for DAQ(AA) and from 0.72 to 0.92 for DAQ(Cα) (Fig. S1). This improvement indicates that differences in DAQ(Cα) primarily arise from spatial discretization, as Cα scores are sensitive to precise atomic positions. We therefore recommend 2 Å sampling for efficient real-time DAQ(AA) evaluation, whereas 1 Å sampling can be used when more precise Cα evaluation is needed, although processing large maps typically requires 30–60 minutes.

### 3.2. Computational time

Fig. 3 evaluates the computational efficiency of DAQplugin using the same 45 EM maps corresponding to the 91 protein chains analyzed in Fig. 2. All calculations were performed on an Intel Xeon E5-1650 v4 CPU (6 cores, 12 threads, 3.60 GHz) without GPU acceleration, representing a typical workstation environment. When a compatible GPU is available, DAQplugin can accelerate 3D-CNN inference using the GPU. For example, on a typical Windows laptop with a GPU, GPU acceleration reduced the computation time by approximately up to 6.5-fold. Figure 3 and Table S1 compare the number of indexed grid points with the computational time for input processing (map loading, normalization, and resampling) and DAQ precomputation (3D-CNN inference). The 3D- CNN inference was the most time-consuming step and strongly correlated with the number of grid points, as expected. Nevertheless, 35 of 45 maps (77.8%) were processed within five minutes. Importantly, precomputation is required only once per EM map, and the resulting compressed NumPy arrays (typically 1–50 MB) can be reused across refinement sessions. Once precomputed, DAQ scores can be accessed in real time during interactive model building and refinement.

**Figure 3.**
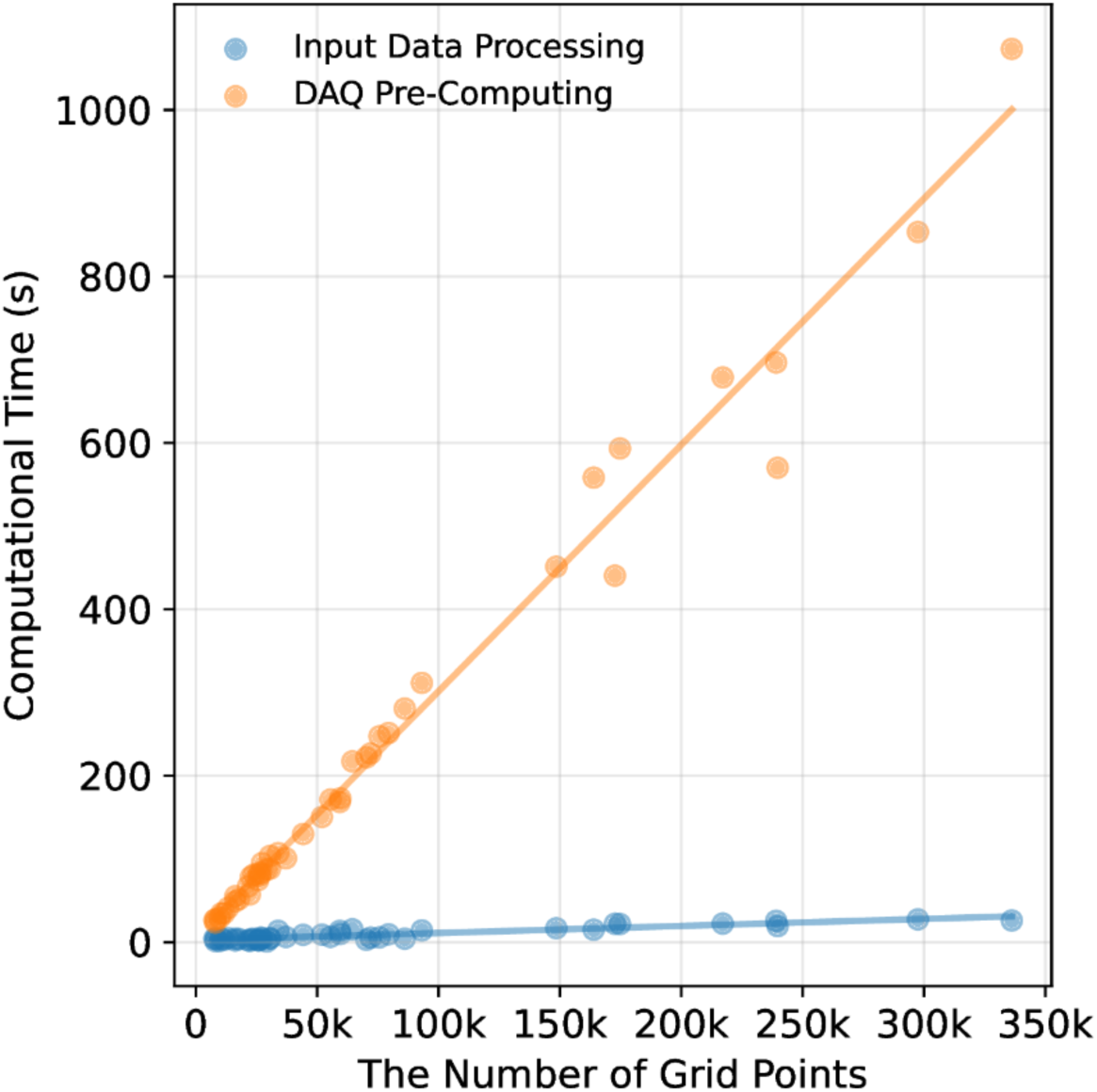
Computational time for DAQplugin. Scatter plots show the computational time for input data processing (blue) and DAQ precomputation by 3D-CNN inference (orange) as a function of the number of grid points. Solid lines indicate linear regression fits for each process.

### 3.3. Case Studies

In this section, we demonstrate how DAQplugin identifies regions requiring refinement and guides interactive model correction.

#### 3.3.1. Correcting sequence register shift

The first example is the mannose transporter complex (PDB ID: 6K1H), a heterohexameric complex^20^. Two model versions (v1-3 and v2-2) are available for this PDB entry. We used version v1-3 together with its corresponding cryo-EM map (EMD-9906; 3.52 Å resolution) as the starting model. The DAQplugin-guided refinement workflow is illustrated in Fig. 4a–e and Movie S2.

**Figure 4.**
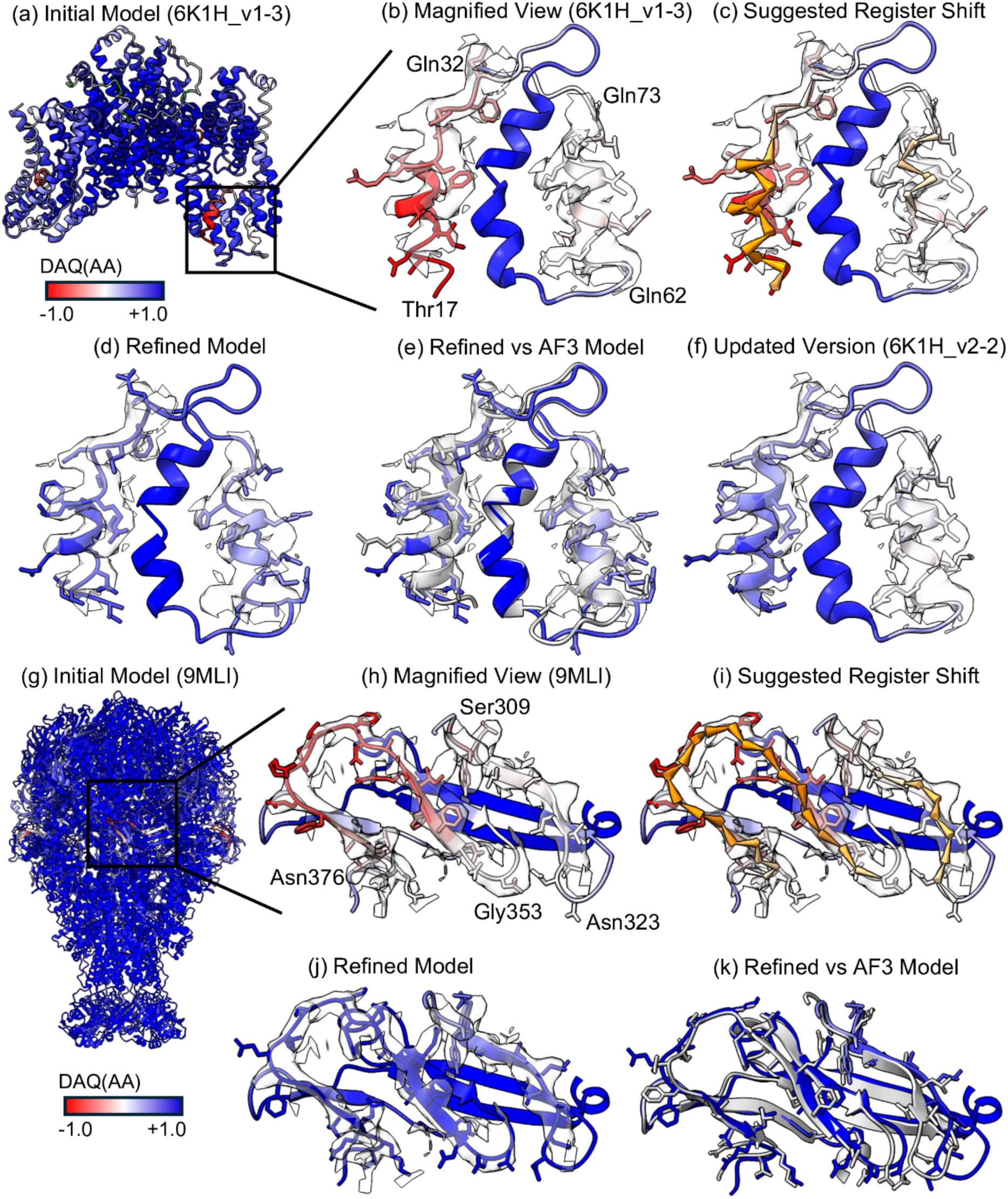
Examples of model validation and sequence register-shift refinement guided by DAQplugin. (a–f) Mannose transporter (EMD-9906, 3.52 Å resolution; PDB ID: 6K1H). **(a)** Initial deposited model (version 1-3). **(b)** Magnified view of two regions in chain C, Thr17–Gln32 and Glu62–Glu73, with low DAQ(AA) scores. Residues are colored by DAQ(AA), from red (negative) through white (0) to blue (positive), with a range of −1.0 to +1.0. **(c)** Sequence register-shift suggestions from DAQplugin. Orange arrows indicate shifts predicted to improve the average DAQ(AA) score by >0.5. **(d)** Refined model after ISOLDE refinement using DAQ-guided positional restraints, followed by correction of side-chain rotamer outliers with DaQRotFix. The model is colored by DAQ(AA). **(e)** Comparison of the refined model with the AlphaFold3 prediction (gray). **(f)** Comparison with the updated deposited model (version 2-2). The Thr17–Gln32 region is consistent with the refined model, whereas Glu62–Glu73 retains the original sequence register. **(g–k)** Insecticidal toxin complex (EMD-48373, 3.4 Å resolution; PDB ID: 9MLI). **(g)** Initial deposited model. **(h)** Magnified view of two regions in chain A, Ser309–Asn323 and Asn354–Asn376, with low DAQ(AA) scores. **(i)** Sequence register-shift suggestions from DAQplugin. **(j)** Refined model after ISOLDE refinement using DAQ-guided positional restraints, followed by correction of side-chain rotamer outliers with DaQRotFix. **(k)** Comparison of the refined model with the AlphaFold3 prediction (gray).

Figure 4a shows the DAQ(AA) scores of the starting model (v1-3), with several regions exhibiting low scores (white or red). A close-up view highlights two problematic regions in chain C, Thr17–Gln32 and Glu62–Glu73, with average DAQ(AA) scores of −0.91 and −0.04, respectively (Fig. 4b). As the −0.5 is strong indicator of incorrect residue placement, the first region would need to be corrected. Using the sequence register-shift suggestion function, DAQplugin indicated that Thr17–Gln32 required a one-residue shift toward the C-terminus, whereas Glu62–Glu73 required a shift toward the N-terminus (Fig. 4c). DAQplugin then generated positional restraints to guide interactive molecular-dynamics-based refinement in ISOLDE, allowing the helices to move toward the suggested positions while maintaining physically realistic geometry (Movie S2). Following register-shift correction, remaining side-chain rotamer outliers were corrected using DaQRotFix (Fig. S2a).

The resulting model is shown in Fig. 4d. After refinement, the average DAQ(AA) score increased from −0.91 to 0.75 for Thr17–Gln32 and from −0.04 to 0.62 for Glu62– Glu73, indicating substantially improved agreement with the local density. The refined model was further compared with an AlphaFold3^21^ (AF3) prediction (Fig. 4e) and the updated deposited structure (v2-2; Fig. 4f). For Thr17–Gln32, both the AF3 and v2-2 models showed the same sequence register as the DAQplugin-refined model, supporting the suggested correction. For Glu62–Glu73, however, the AF3 model agreed with the DAQplugin-refined register, whereas the v2-2 model retained the original register. This region in v2-2 retained a low average DAQ(AA) score of 0.04, compared with 0.62 after DAQplugin-guided refinement. Therefore, the improved DAQ(AA) score and agreement with the AF3 model suggest that the Glu62–Glu73 region in the v2-2 model in the PDB entry may still require revision.

The second example (Fig. 4g–k) demonstrates correction of sequence register shifts in β-strands. The target is a subunit of an insecticidal toxin complex (EMD-48373, 3.4 Å resolution; PDB ID: 9MLI), which forms a homo-pentamer containing a total of 12,595 residues^22^. Despite the large size of the complex, DAQplugin identified two small regions with low DAQ(AA) scores, highlighting its ability to detect localized modeling errors that may otherwise be difficult to recognize. Among the five chains, these regions were found only in chain A, within the receptor-binding domain (RBD): Ser309–Asn323 and Gly353– Asn376, with average DAQ(AA) scores of −0.09 and −0.57, respectively. DAQplugin predicted that shifting both regions by one residue toward the N-terminus would improve their DAQ(AA) scores (Fig. 4i). The refinement trajectory is shown in Movie S3.

Using DAQplugin-generated positional restraints, interactive refinement was performed in ISOLDE to shift the sequence assignment along the β-strands while maintaining realistic backbone geometry and β-sheet hydrogen bonding. Because a one-residue register shift in a β-strand also requires reversal of side-chain orientations, the nonbonded interaction scaling factor, λ(equil), was initially reduced to 0.1 to facilitate side-chain rearrangement. Remaining rotamer outliers after refinement were corrected using DaQRotFix (Fig. S2b). In the refined model (Fig. 4j), the average DAQ(AA) scores increased to 0.74 and 1.20 for Ser309–Asn323 and Gly353–Asn376, respectively.

Unlike the first example, PDB entry 9MLI has no revised version for comparison. Instead, we compared the structures with an AlphaFold3 prediction. The refined model showed substantially better agreement with the AlphaFold3 structure, with the Cα RMSD decreasing from 3.44 Å to 1.36 Å (Fig. 4k). Together with the improved DAQ(AA) scores, this agreement supports the proposed register shifts and suggests that these regions in the deposited PDB model may contain sequence-register errors.

#### 3.3.2. Global positional shift correction

The next example (Fig. 5) is the tetrameric secretory immunoglobulin A complex with polymeric immunoglobulin receptor and joining chain (EMD-20751, 2.9 Å resolution; PDB ID: 6UE9)^23^. Movie S4 shows real-time updates of DAQ(Cα) during refinement. The deposited model showed reasonable overall agreement with the cryo-EM map, with a cross-correlation coefficient of 0.64. However, DAQ(AA) and DAQ(Cα) revealed distinct patterns. DAQ(AA) was predominantly positive (mean 0.37; Fig. 5a), whereas DAQ(Cα) was broadly negative (mean −0.44; Fig. 5b). This suggests that the residue assignments were generally consistent with the local density, but the Cα positions were systematically displaced from their optimal locations. We therefore performed rigid-body fitting of the model into the cryo-EM map using the *fitmap* command in ChimeraX. The fitted model was displaced by 1.9 Å RMSD from the deposited structure. Following fitting, the mean DAQ(AA) increased from 0.37 to 0.63, while DAQ(Cα) improved substantially from −0.44 to 0.53 (Fig. 5c, 5d), indicating improved model–map agreement.

**Figure 5.**
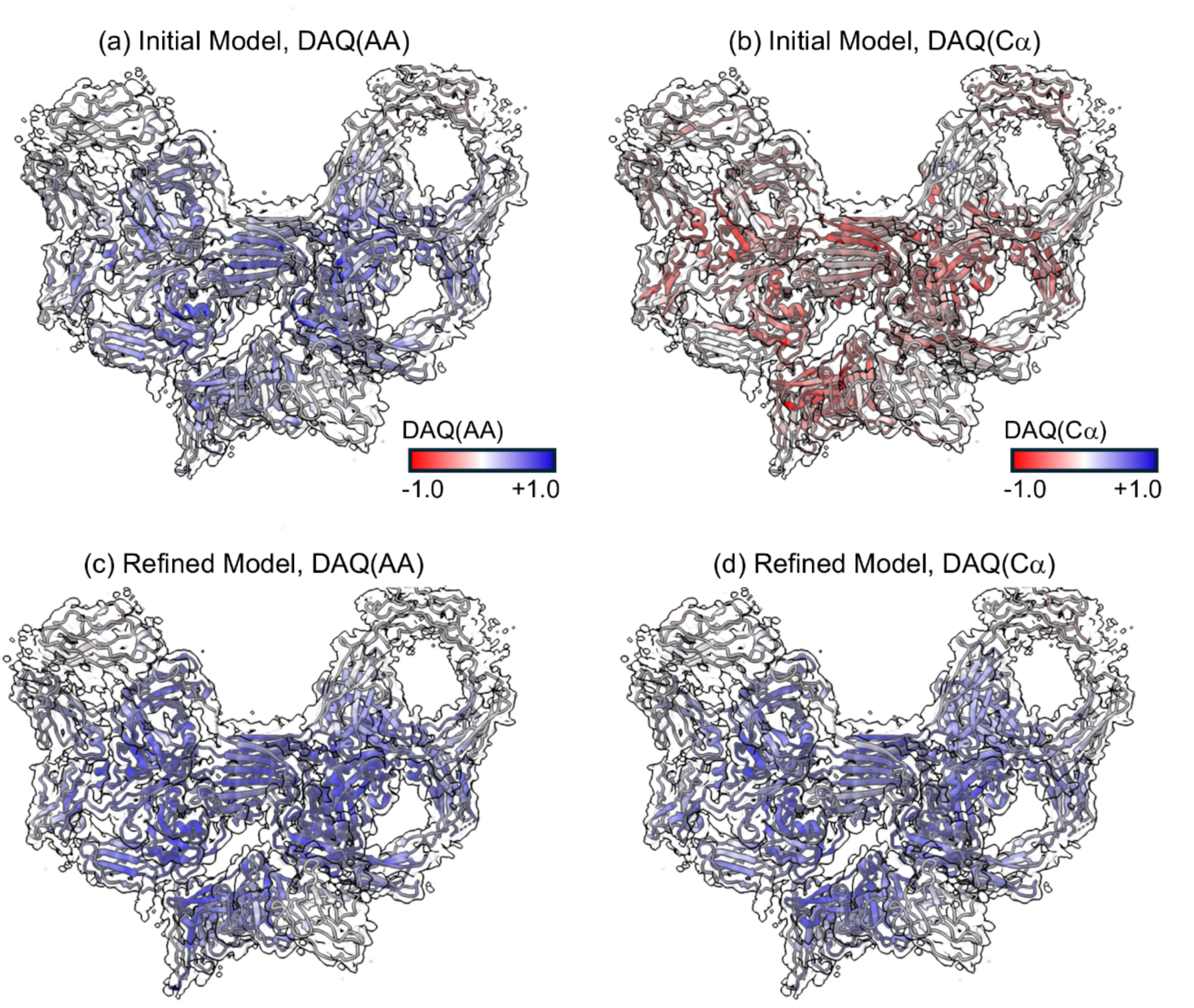
Positional shift correction guided by DAQ(Cα). The example shows tetrameric secretory immunoglobulin A with polymeric immunoglobulin receptor and joining chain (EMD-20751, 2.9 Å resolution; PDB ID: 6UE9). **(a)** Initial deposited model colored by DAQ(AA), showing generally positive scores. **(b)** Initial model colored by DAQ(Cα), revealing broadly negative scores indicative of positional inconsistency. **(c)** Model after rigid-body fitting to the cryo-EM map using the ChimeraX *fitmap* command, colored by DAQ(AA). The mean DAQ(AA) improved from 0.37 to 0.63. **(d)** Fitted model colored by DAQ(Cα), with the mean score improving from −0.44 to 0.53.

Importantly, the original model had a relatively high cross-correlation coefficient of 0.64, which did not readily reveal the positional error. In contrast, DAQ(Cα) directly highlighted the systematic backbone displacement through broadly negative residue-wise scores. This example illustrates how DAQ(Cα) can complement global map–model correlation by identifying positional errors that may remain difficult to recognize from global agreement metrics alone.

#### 3.3.3. Model assessment and refinement assisted by DAQplugin

Beyond sequence register and global shift correction discussed above, DAQplugin can assist more general model assessment and refinement by identifying regions with poor model–map agreement and providing real-time feedback during correction. We demonstrate these applications using two examples.

The first example is SLC40/ferroportin in complex with Fab (EMD-21460, 3.0 Å resolution; PDB ID: 6VYH)^24^. The initially deposited model (Fig. 6a, b) showed multiple regions with negative DAQ(AA) scores across the C-terminal domain of chain C (Ala129– Ala233), with an average score of −0.10. To refine this domain, we used an AlphaFold3 (AF3) model of the light chain as structural guidance. After superposition onto the initial structure, the N-terminal domain (Asp21–Glu128) aligned well, whereas the C-terminal domain adopted a substantially different orientation. We therefore isolated the C-terminal domain from the AF3 model, manually positioned it into the cryo-EM density (an intermediate step is shown in Fig. 6c), and further fitted it using the ChimeraX *fitmap* command, followed by refinement in ISOLDE.

**Figure 6.**
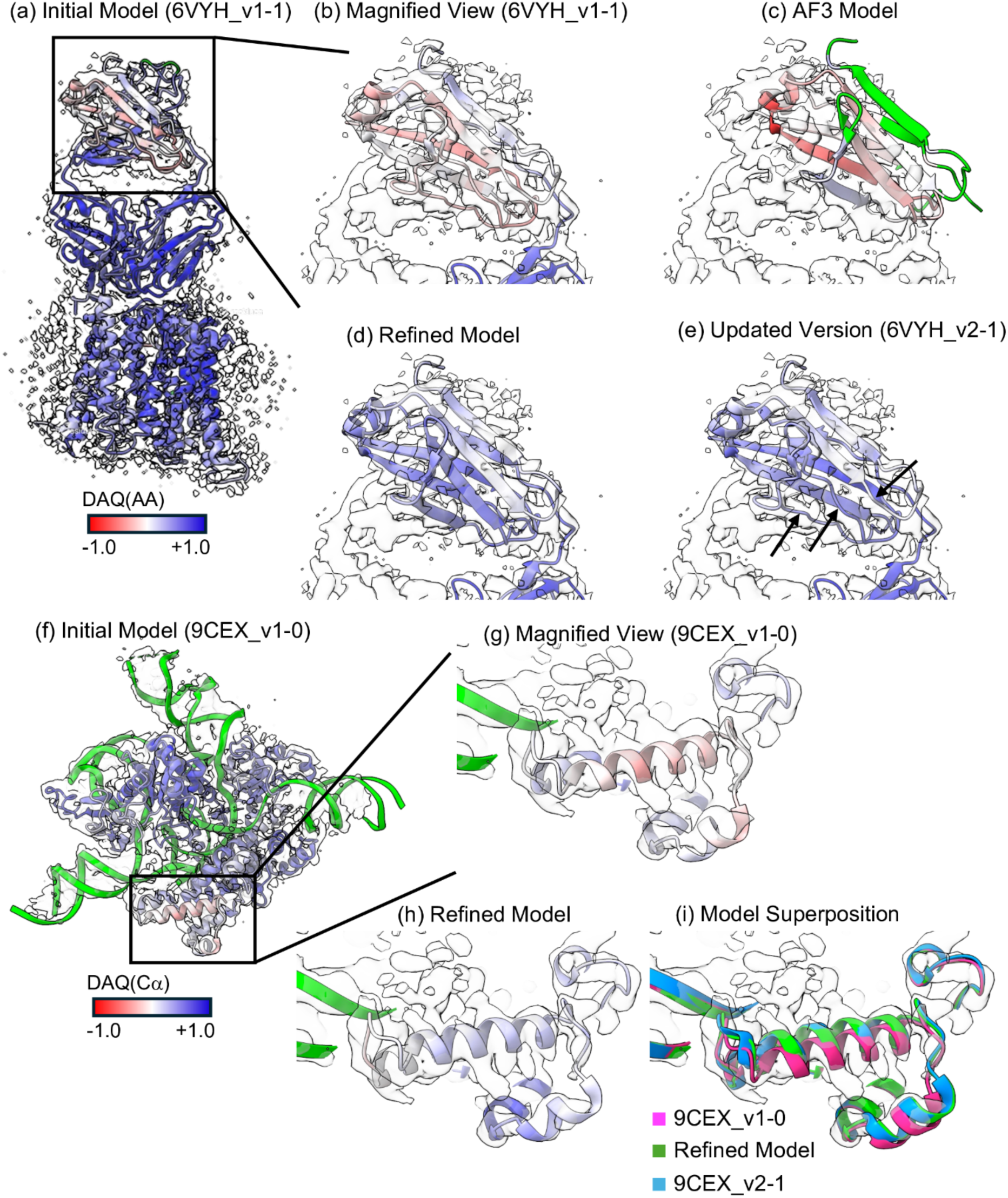
Examples of model assessment and refinement using DAQplugin. Residues evaluated by DAQ are colored from red (negative) through white (0) to blue (positive), with a range of −1.0 to +1.0. Protein residues outside the region above half of the user-defined contour level, as well as DNA/RNA residues, are not evaluated by DAQplugin. These unevaluated regions are shown in green. **(a–e)** SLC40/ferroportin in complex with Fab (EMD-21460, 3.0 Å resolution; PDB ID: 6VYH). **(a)** Initial deposited model (version 1-1) colored by DAQ(AA). **(b)** Magnified view of the light-chain C-terminal domain. **(c)** An intermediate snapshot during manual repositioning of the C-terminal domain of AF3 model into the cryo-EM density. DAQ(AA) scores are updated in real time as the domain is moved. The domain was subsequently positioned by manual adjustment followed by fitting with the ChimeraX *fitmap* command. **(d)** The refined model after domain replacement and subsequent ISOLDE refinement. The average DAQ(AA) score for Ala120–Ala233 improved from −0.10 to 0.47. **(e)** Updated deposited model (version 2-1). The refined model showed closer agreement with the updated structure, with a Cα RMSD of 0.67 Å compared with 6.28 Å between the initial and updated models. Three arrows highlight the irregular β-strand regions observed in the updated PDB model. **(f–i)** the Spizellomyces punctatus Fanzor (SpuFz) State 4 (EMD-45522, 3.27 Å resolution; PDB ID: 9CEX). **(f)** Initial deposited model (version 1-0) colored by DAQ(Cα). RNA is colored in green. **(g)** Magnified view of two α-helices spanning Leu171–Asp205 with negative DAQ(Cα) scores. **(h)** The refined model after local ISOLDE refinement of this region. The average DAQ(Cα) score improved from −0.10 to 0.17. **(i)** Superposition of the initial v1-0 model (magenta), the refined model (green), and the updated v2-1 model (blue).

After refinement, the average DAQ(AA) score for Ala129–Ala233 improved from −0.10 to 0.47 (Fig. 6d). The refined model closely agreed with the subsequently updated deposited structure (PDB 6VYH version 2-1; Fig. 6e), with a Cα RMSD of 0.67 Å, compared with 6.28 Å between the initial (v1-1) and updated (v2-1) PDB models. The differences between our refined and v2-1 models were primarily localized to loop and irregular β-strand regions observed in the updated PDB model (Fig. 6e). Both models also showed similar positive average DAQ(AA) scores (0.47 for our refined model and 0.45 for v2-1), indicating comparable agreement with the cryo-EM density. Although an AF3 prediction is not necessarily more accurate than an experimentally derived model, in this case its C-terminal domain provided useful structural guidance for correcting the region identified by DAQplugin.

The next example is *Spizellomyces punctatus* Fanzor (SpuFz) State 4 (EMD-45522, 3.27 Å resolution; PDB ID: 9CEX)^25^. In the initial deposited PDB model (version 1-0), two α-helices spanning Leu171–Asp205 showed negative DAQ(Cα) scores, with an average of −0.10, indicating a region requiring further inspection and refinement (Fig. 6f, g). This region was interactively refined by selecting these regions and running ISOLDE while monitoring DAQ(Cα) scores in real time. After refinement (Fig. 6h), the average DAQ(Cα) increased to 0.17, and the resulting structure became more similar to the subsequently updated PDB model (version 2-1) (Fig. 6i). Compared with the initial version model (magenta) in Fig. 6i, the refinement shifted the long helix (res. Arg178 to Leu193) upwards in the figure and the small helix (res. Glu200 to Asp205) toward the right, making the refined model (green) closely agree with the v2-1 model (cyan). The Cα RMSD for Leu171–Asp205 decreased from 0.86 Å between the initial and updated models to only 0.50 Å between the DAQ-guided refined and updated models, with the remaining structural differences being essentially indistinguishable by model validation and map–model agreement metrics.

## 4. Discussion

We developed DAQplugin for real-time deep-learning-based validation of protein models against cryo-EM maps during interactive modeling in ChimeraX. Unlike conventional post- modeling validation, DAQplugin updates residue-wise model–map agreement as structures are modified, integrating validation directly into the modeling and refinement process.

A key challenge for real-time validation is the computational cost of 3D-CNN inference. DAQplugin addresses this by precomputing DAQ score grids once per cryo-EM map and using rapid KD-tree-based lookup during refinement. This enables real-time score updates on standard CPUs without GPU acceleration.

Several limitations remain. Although precomputation enables real-time DAQ scoring, processing very large cryo-EM maps can still be time-consuming. The current implementation runs on standard CPUs to maximize accessibility and simplify installation on standard workstations and laptops. When a compatible GPU is available, GPU acceleration can reduce the precomputation time. Nevertheless, processing large cryo-EM maps may still require several minutes or longer, depending on the map size. Other limitations originate from the current DAQ framework itself. DAQ evaluates protein models primarily at the residue and backbone levels, including amino-acid identity and Cα positions, but does not explicitly evaluate side-chain conformations and currently does not support nucleic acids or ligands.

Future development will extend DAQ to these additional structural components and modeling environments. We plan to develop support for Coot^11^, which will require adapting the current interactive scoring and model-update framework to its distinct software architecture and interfaces. We also envision closer integration with AI-based structure modeling and refinement methods^26–28^, including AlphaFold3^21^, allowing predicted structures and cryo-EM density to be evaluated and refined together within an interactive modeling workflow.

## Acknowledgements

This work was partly supported by the National Institutes of Health (R01GM133840, R35GM158267, R21AI187928) to DK and the National Science Foundation (DBI2433490 to GT, IIS2211598, DBI2146026, and DBI2422620 to DK).

## Data availability

DAQplugin is available in ChimeraX Toolshed (https://cxtoolshed.rbvi.ucsf.edu/apps/chimeraxdaqplugin) and Github repository at https://github.com/kiharalab/DAQplugin. DaQRotFix is available in ChimeraX Toolshed (https://cxtoolshed.rbvi.ucsf.edu/apps/chimeraxdaqrotfix) and Github repository at https://github.com/kiharalab/DaQRotFix.

## Supporting information

**Table S1.**
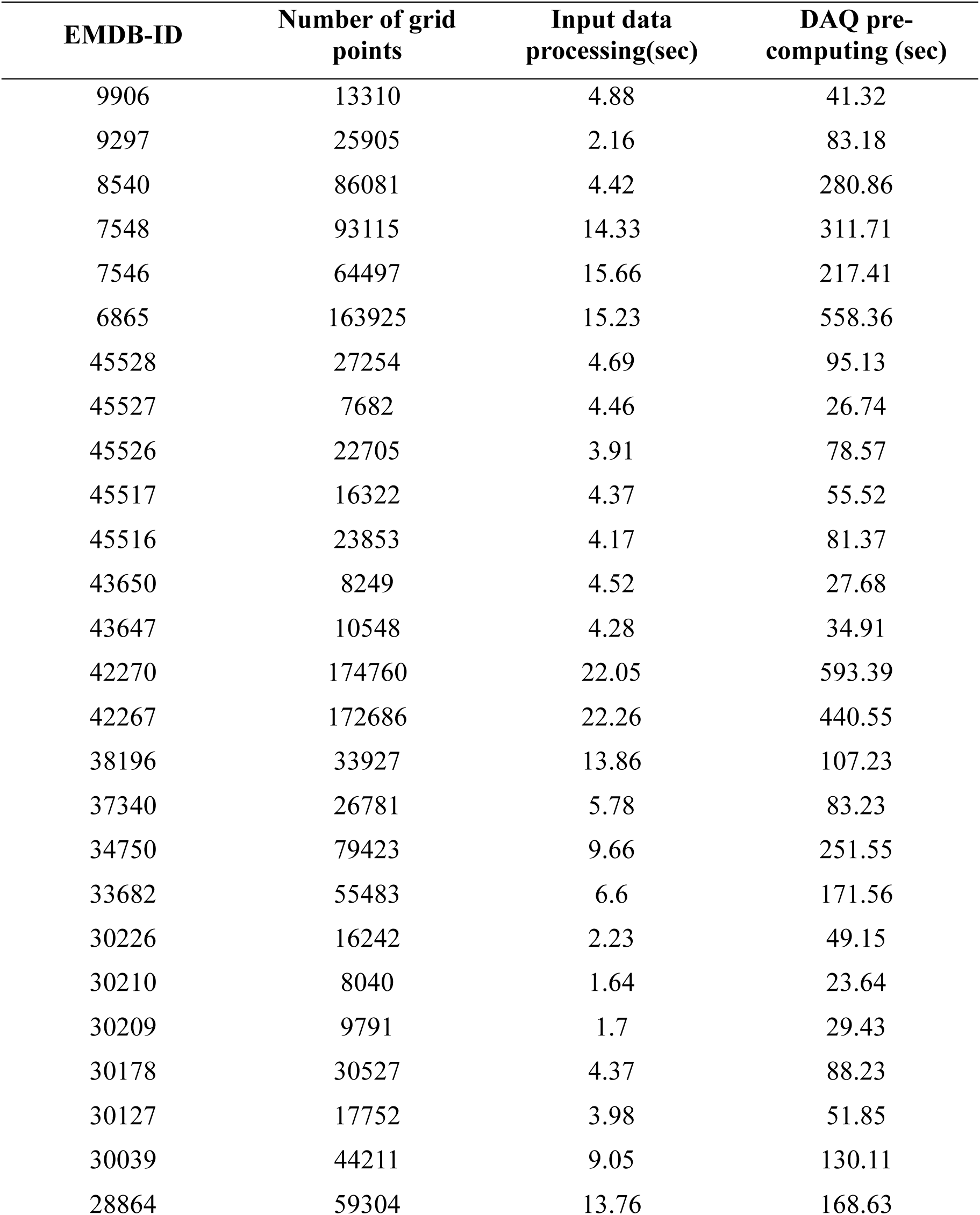
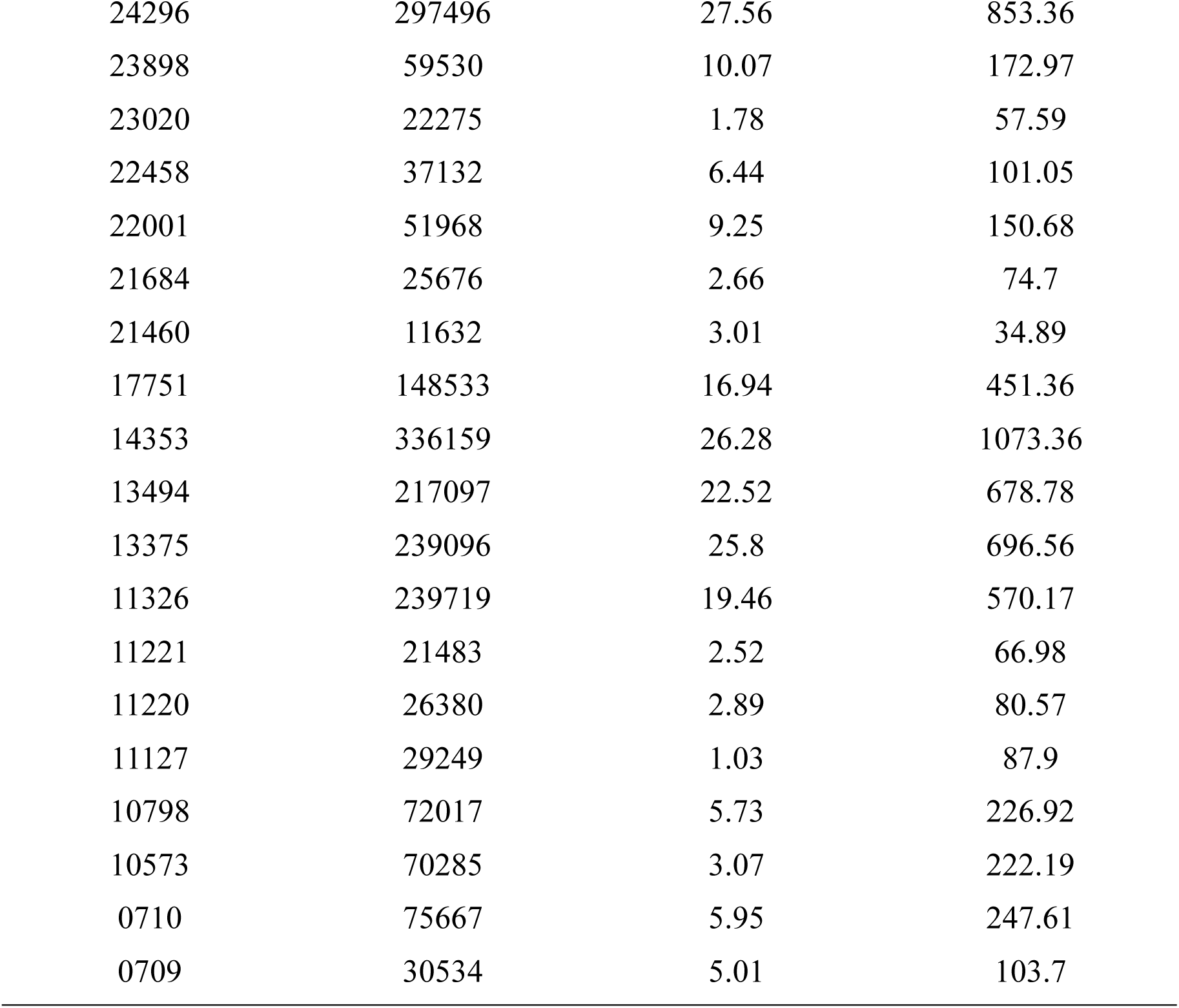
Computational time for DAQplugin pre-computation. Data for 45 EM maps of different sizes were shown. For each map, the number of grid points, input data processing time, and DAQ pre-computing time (3D-CNN inference) are reported.

**Figure S1.**
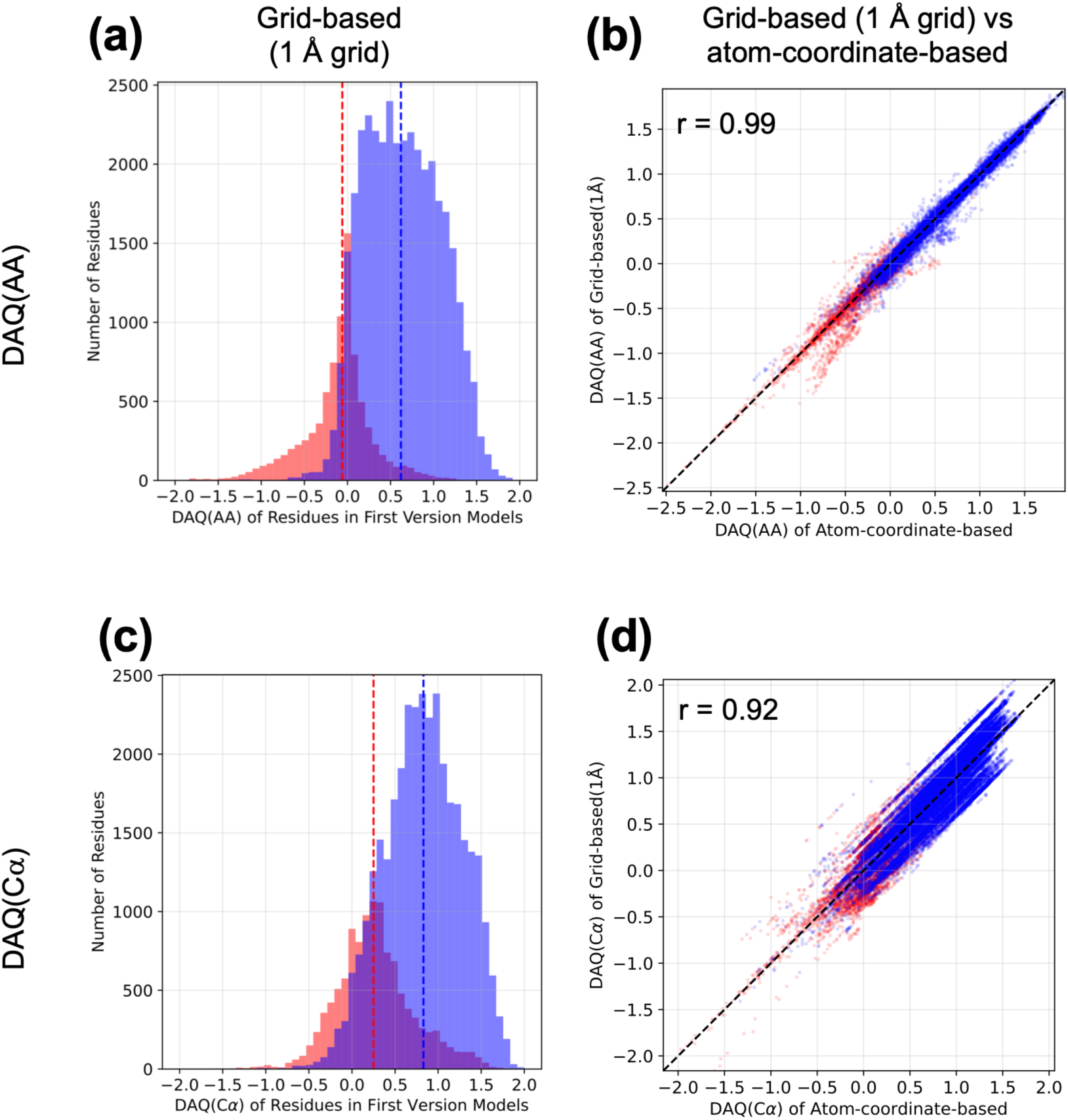
Comparison of 1 Å grid-based and atom-coordinate-based DAQ scores. Dashed vertical lines indicate median values. **(a)** Distribution of 1 Å grid-based DAQ(AA) scores for modified (>2.0 Å Cα displacement) and preserved residues, with median values of −0.06 and 0.62, respectively. **(b)** Residue-wise comparison of 1 Å grid-based and atom-coordinate-based DAQ(AA) scores (Pearson *r* = 0.99). **(c)** Distribution of 1 Å grid-based DAQ(Cα) scores for modified (>1.0 Å Cα displacement) and preserved residues, with median values of 0.25 and 0.83, respectively. **(d)** Residue-wise comparison of 1 Å grid-based and atom-coordinate-based DAQ(Cα) scores (Pearson *r* = 0.92).

**Figure S2.**
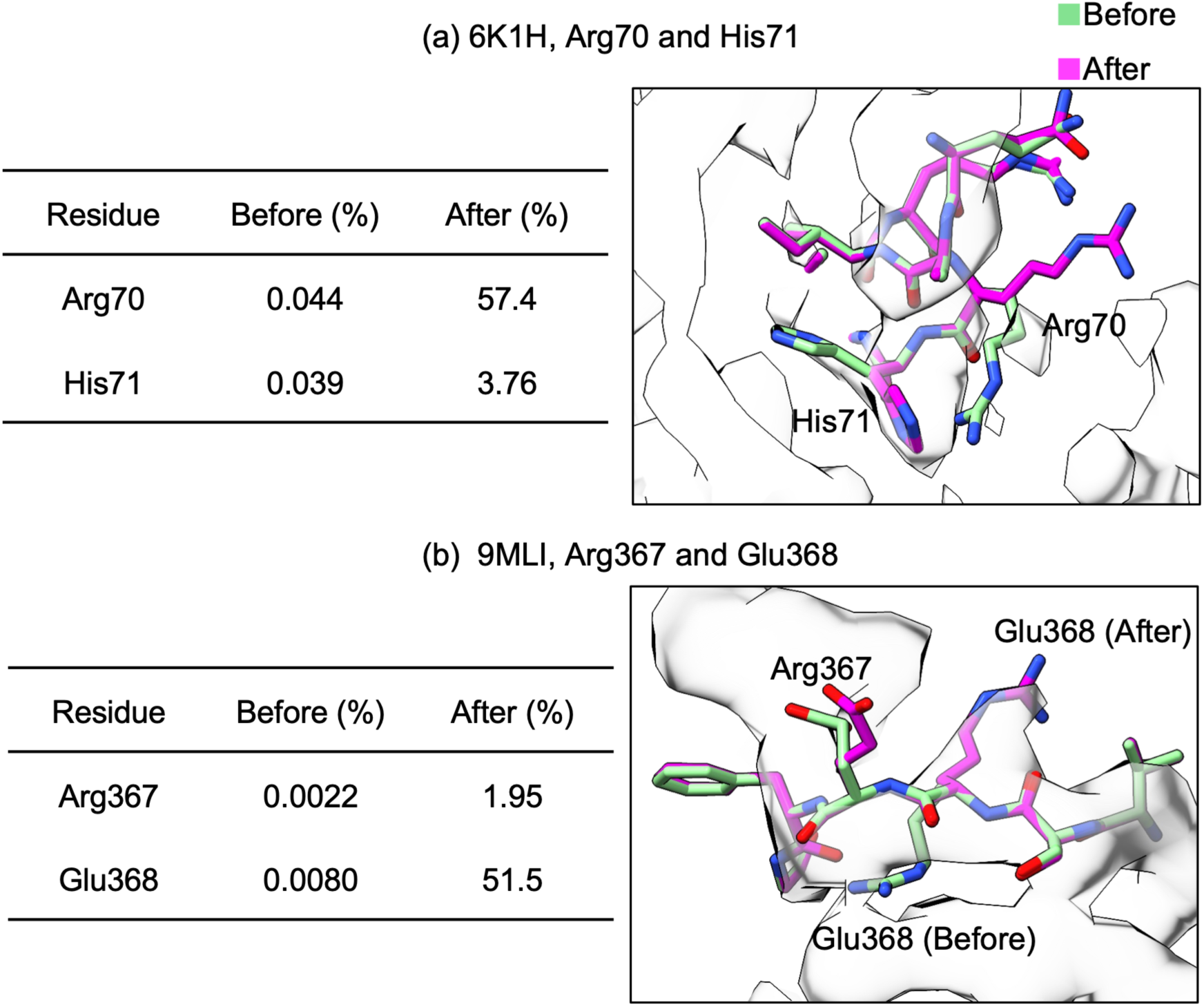
Examples of side-chain rotamer outlier correction using DaQRotFix. Structures before and after correction are shown in green and magenta, respectively, and the tables show the corresponding rotamer probabilities. The initial structures were obtained after DAQ-guided register-shift refinement in ISOLDE and contained rotamer outliers with probabilities below 0.05%. Cryo-EM densities are shown as gray surfaces, contoured at 0.12 for EMD-9906 and 0.23 for EMD-48373. **(a)** Arg70 and His71 in PDB 6K1H_v1-3 and EMD-9906 before and after correction. DaQRotFix increased the rotamer probabilities from 0.044% to 57.4% for Arg70 and from 0.039% to 3.76% for His71. **(b)** Arg367 and Glu368 in PDB 9MLI and EMD-48373 before and after correction. DaQRotFix increased the rotamer probabilities from 0.0022% to 1.95% for Arg367 and from 0.0080% to 51.5% for Glu368.

**These supplementary Movies are provided in separate files (MP4 format)**

**Supplementary Movie S1. Interactive refinement of a sequence register error using DAQplugin (7FET).** The example shows PDB ID 7FET, chain B, with the corresponding cryo-EM map EMD- 31563, and corresponds to the workflow shown in Fig. 1b, panels 1–4. DAQplugin identified a region with negative DAQ(AA) scores (Gly593–Pro621; average DAQ(AA), −0.51) and indicated the suggested direction of the sequence register shift with orange arrows. The region was then interactively refined in ISOLDE following the suggested direction. During refinement, DAQ(AA) scores were continuously updated and visualized on the structure in real time. After refinement, the sequence register was corrected, and the average DAQ(AA) score increased from −0.51 to 0.89.

**Supplementary Movie S2. Interactive refinement of sequence register errors in two helical regions using DAQplugin (6K1H).** The example shows two regions, Thr17–Gln32 and Glu62–Glu73, in chain C of PDB ID 6K1H (version 1-3), with the corresponding cryo-EM map EMD-9906. The movie corresponds to the refinement shown in Fig. 4a–d. DAQplugin identified regions with low DAQ(AA) scores and provided guidance for correcting their sequence registers. The model was interactively refined in ISOLDE while DAQ(AA) scores were continuously updated and visualized on the structure in real time. After refinement, the average DAQ(AA) score increased from −0.91 to 0.75 for Thr17–Gln32 and from −0.04 to 0.62 for Glu62–Glu73.

**Supplementary Movie S3. Interactive refinement of sequence register errors in two β- strand regions using DAQplugin (9MLI).** The example shows two regions, Ser309–Asn323 and Gly353–Asn376, in chain A of PDB ID 9MLI, with the corresponding cryo-EM map EMD-48373. The movie corresponds to the refinement shown in Fig. 4g–j. DAQplugin identified the two regions with negative average DAQ(AA) scores of −0.09 and −0.57, respectively, and suggested shifting both regions by one residue toward the N-terminus. The regions were then interactively refined in ISOLDE while DAQ(AA) scores were continuously updated and visualized on the structure in real time.

**Supplementary Movie S4. Interactive correction of Cα positional shifts using DAQplugin (6UE9).** The example shows PDB ID 6UE9 with the corresponding cryo-EM map EMD-20751 and corresponds to the refinement shown in Fig. 5. In the initial model, DAQ(Cα) scores were broadly negative, with a mean of −0.44, indicating systematic displacement of Cα positions relative to the local density. The model was rigid-body fitted into the map using the ChimeraX *fitmap* command, while DAQ(Cα) scores were continuously updated and visualized on the structure in real time. Following fitting, the mean DAQ(Cα) score increased from −0.44 to 0.53.

